# Developing And Internally Validating AI-Based Aging Resilience Biomarkers in Non-Human Primates

**DOI:** 10.64898/2026.02.25.707531

**Authors:** Robert F. Bennett, Jaime L. Speiser, John D. Olson, George W. Schaaf, Thomas C. Register, J. Mark Cline, Laura A. Cox, Ellen E. Quillen

**Affiliations:** Wake Forest School of Medicine

## Abstract

Quantifying biological aging is crucial for understanding functional decline before the onset of morbidity. While many accelerated aging and frailty measures based on clinical data exist for humans and several for rodent models of aging, there are few options for non-human primates (NHPs). NHP clinical data has several unique features including a lack of clinically delineated normative values for features and variability in data collection over long lifespans. There are also wide discrepancies in the number of available clinical measures and number of animals across data sets. To address these challenges, we developed and validated “Aging Resilience” (AR) metrics using longitudinal, routine clinical data from two distinct non-human primate cohorts: 4,328 baboons and 281 rhesus macaques. We trained five computational models—including Linear Mixed-Effects Models, Random Forest, and Recurrent Neural Networks (RNN)—to predict chronological age, subsequently deriving AR metrics that represent the velocity (Rate of Aging) and cumulative burden (Normalized Cumulative Aging) of physiological deviation. While linear models achieved high precision in predicting chronological age (test R^2^ up to 0.99), they correlated poorly with actual lifespan. In contrast, AR metrics derived from non-linear models (RNN and Random Forest) displayed strong predictive validity for mortality (Pearson’s r > 0.8). These findings highlight a critical paradox: models that best predict chronological age do not necessarily capture the biological resilience determining healthspan. This study establishes a scalable framework for monitoring biological aging in translational models using standard veterinary records.

## Introduction

The quantification of biological aging is a crucial goal in geroscience, offering the potential to move beyond chronological age to understand an individual’s functional decline and mortality risk —often driven by the systemic physiological dysregulation described collectively as ‘Hallmarks of Aging’— before the onset of morbidities^1^. While numerous biomarkers of aging have been proposed, many lack robust validation or require specialized assays (e.g., omic data analysis) not available in standard clinical practice^2^. Large, longitudinal (repeated measures) datasets of routine clinical measurements, however, present an underutilized resource for developing more holistic and dynamic aging clocks. This is especially critical because cross-sectional biomarkers capture only a static snapshot of accumulated damage, whereas longitudinal data allow for an understanding of change over time. The primary challenge lies in effectively modeling complex and time-dependent relationships within this high-dimensional data to produce a generalizable predictor of age.

Machine learning and other advanced computational methods are well-suited for this challenge, capable of modeling subtle patterns in longitudinal data that traditional statistical models may miss^3^. The development of an accurate metric that represents the burden of aging can serve two purposes: first, as a means of extracting relevant features predicting health status, and second, as a foundation for deriving unique biomarkers of biological health status, which we collectively refer to as Aging Resilience (AR) metrics. The deviation of predicted age from chronological age over time may represent an individual’s AR, a concept which could prove highly predictive of lifespan and healthspan. Within this framework, we distinguish between two methodological approaches: Rate of Aging (RoA), which estimates the slope of these deviations to capture the velocity of aging, and Normalized Cumulative Aging (NCA), which quantifies the total burden of deviation over time relative to the observation period.

This distinction is uniquely relevant for non-human primates (NHPs) because humane euthanasia is often indicated at the first onset of serious illness or visible health decline, diminishing the usefulness of the types of frailty/multi-morbidity indices used in humans^4^. Therefore, sensitive metrics like AR—whether calculated as RoA or NCA—which can quantify an individual’s resilience level compared to chronological aging from routine data long before overt morbidity, are essential for advancing geroscience in these critical translational models.

To date, no such resilience metric exists for NHPs. While the deficit accumulation model—exemplified by frailty indices generated for both human and rodent models^5–7^—is a cornerstone of geroscience, its application in NHPs is complicated by a lack of standardized clinical reference ranges and cutoffs necessary to define specific deficits^8^. Moreover, traditional frailty assessments typically identify decline only after functional deficits have accumulated. In contrast, the AR approach proposed here utilizes continuous deviations in routine biomarkers to quantify resilience prior to the onset of serious health decline or disease. Although similar computational strategies have been successfully explored in murine models^9^, a comparable framework has yet to be established for primates. Bridging this gap is essential, as the NHP offers a uniquely high-fidelity translational model for testing metrics intended for eventual human implementation. By utilizing clinical data structures analogous to human electronic health records, these cohorts provide the high-quality, longitudinal input necessary to overcome the data limitations that have historically hindered the application of AI in precision medicine^10^.

In this study, we systematically developed and internally validated six distinct AI models for the prediction of chronological age using longitudinal clinical data from two separate non-human primate cohorts with contrasting numbers of observations and features. We evaluated the performance of Multiple Linear Regression (MLR), Linear Mixed-Effects Models (LMM), Random Forest (RF), a feed-forward Neural Network (NN), and a Recurrent Neural Network (RNN). Subsequently, we derived AR metrics (RoA and NCA) from the best-performing models and assessed their validity by correlating them with the known lifespan of the subjects. The goal of this research was to develop a workflow that generates the most biologically informative representation for AR using these two distinctly different longitudinal NHP cohorts, creating a metric that can further our understanding of biological aging.

## Methods

This study used longitudinal, repeated measures of lab, clinical, and imaging data (listed in Supplementary Materials) from two distinct non-human primate cohorts (Table 1). The first, from the Southwest National Primate Research Center (SNPRC), is a large dataset comprising 9,836 total observations (total number of ID/Year combinations, after collapsing by Year) across 4,328 baboons *(Papio spp*.), with a nearly 2:1 female-to-male ratio. The ages in this cohort range from 6 to 33 years (roughly equivalent to 18–99 human years) and include 19 clinical features. The second cohort, the Radiation Late Effects Cohort (RLEC), is smaller and more feature rich. It contains 1,227 observations from 281 rhesus macaques *(Macaca mulatta)*, with a female-to-male ratio of nearly 1:3. The RLEC data spans ages 6 to 24 (roughly equivalent to 18–72 human years) and contains a total of 80 clinical features. The notable differences in sample size, sex distribution, and feature depth between the two cohorts provide a robust basis for evaluating the generalizability of our modeling approaches.

**Table 1.** Data overview of both cohorts – the SNPRC data contains a total of 19 features and RLEC contains 80 – Average Age is across all timepoints.

| Cohort | Sex | Total Obs. | Unique Animals | Min. Age | Max. Age | Average Age |
| --- | --- | --- | --- | --- | --- | --- |
| SNPRC | Female | 6,604 | 2,723 | 6 | 33 | 12.615 |
| SNPRC | Male | 3,232 | 1,605 | 6 | 27 | 10.339 |
| **SNPRC** | **Total** | **9,836** | **4,328** | **6** | **33** | **11.867** |
| RLEC | Female | 276 | 74 | 6 | 20 | 10.549 |
| RLEC | Male | 951 | 188 | 6 | 24 | 9.069 |
| **RLEC** | **Total** | **1,227** | **281** | **6** | **24** | **9.005** |

**Table 2.** Performance overview of the analyses in SNPRC.

| Model | Variant | RMSE Train | RMSE Val | RMSE Test | R2 Train | R2 Val | R2 Test |
| --- | --- | --- | --- | --- | --- | --- | --- |
| LMM | lmm_random_intercept_bobyqa | 1.18 | 4.944 | 5.117 | 0.95 | 0.126 | 0.159 |
| MLR | with_relTime | 4.137 | 4.444 | 4.522 | 0.356 | 0.324 | 0.344 |
| NN | with_seq_idx | 4.567 | 4.726 | 5.05 | 0.231 | 0.228 | 0.181 |
| RF | with_relTime | 3.478 | 4.322 | 4.485 | 0.569 | 0.36 | 0.353 |
| RNN | base | 4.53 | 4.906 | 5.066 | 0.234 | 0.185 | 0.177 |

**Table 3.** Performance overview of the analyses in RLEC.

| Model | Variant | RMSE Train | RMSE Val | RMSE Test | R2 Train | R2 Val | R2 Test |
| --- | --- | --- | --- | --- | --- | --- | --- |
| LMM | lmm_randomslope_time_bobyqa | 0.254 | 0.297 | 0.242 | 0.995 | 0.994 | 0.992 |
| MLR | base | 0.903 | 1.126 | 0.987 | 0.943 | 0.883 | 0.925 |
| NN | with_relTime | 1.76 | 1.901 | 1.525 | 0.795 | 0.671 | 0.794 |
| RF | with_relTime | 2.165 | 2.336 | 2.184 | 0.78 | 0.506 | 0.628 |
| RNN | base | 1.873 | 1.977 | 1.96 | 0.831 | 0.692 | 0.686 |

**Table 4.**
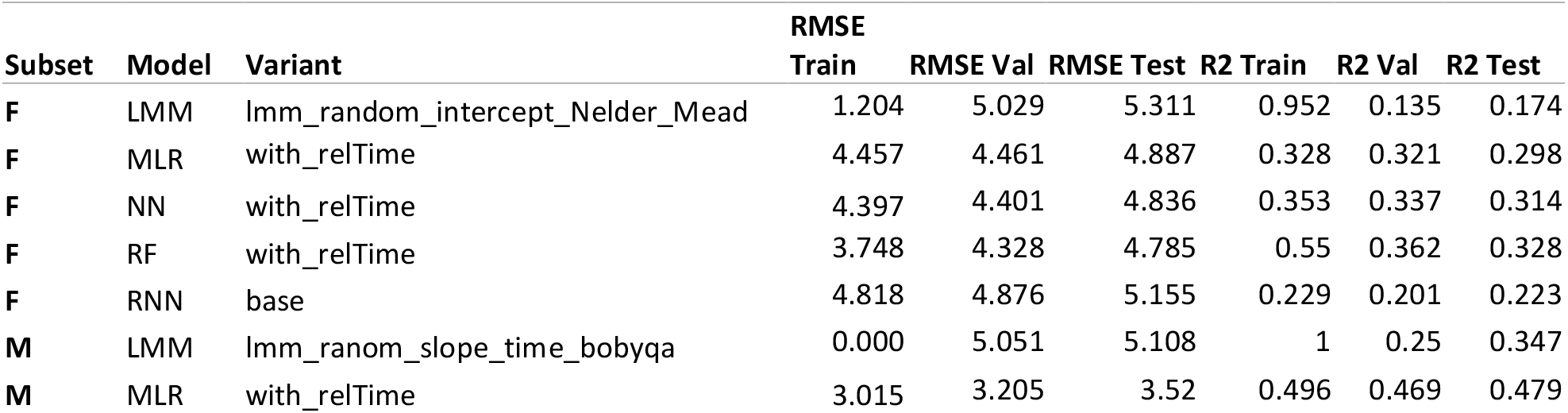
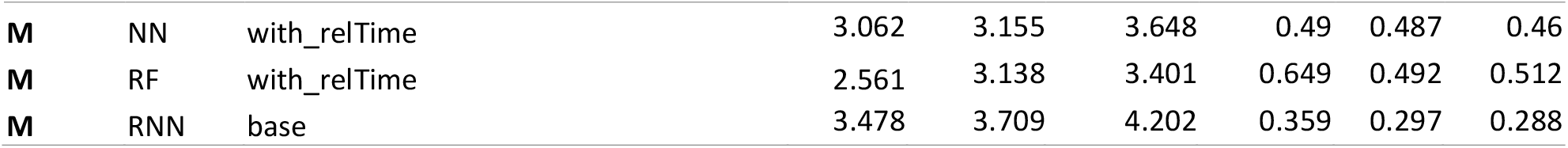
Results overview for the sex-specific analyses in the SNPRC dataset; blue is male and orange is female.

**Table 5.**
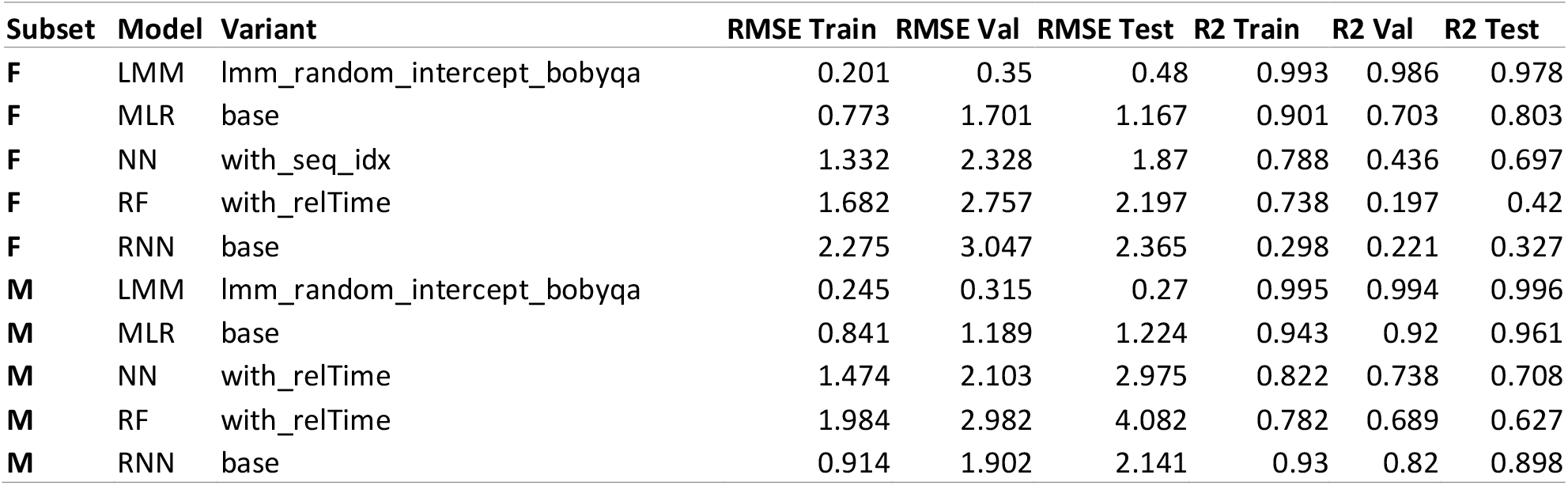
Results overview for the sex-specific analyses in the RLEC dataset; blue is male and orange is female.

We sought to calculate a metric that captures longitudinal aging trajectories and to extract biologically relevant features by training and evaluating six models on the two independent datasets, RLEC and SNPRC. The entire computational workflow, depicted in Figure 2, was implemented in R Statistical Software (v4.4.3; R Core Team, 2024). Data manipulation and visualization were performed using the tidyverse suite (v2.0.0)^11^, while missing values were addressed using the mice package (v3.17.0)^12^. While portions of the pipeline were executed locally, the computationally intensive steps (imputation and model training/testing) were conducted on a Linux-based high-performance computer cluster at Wake Forest University School of Medicine.

**Figure 1.**
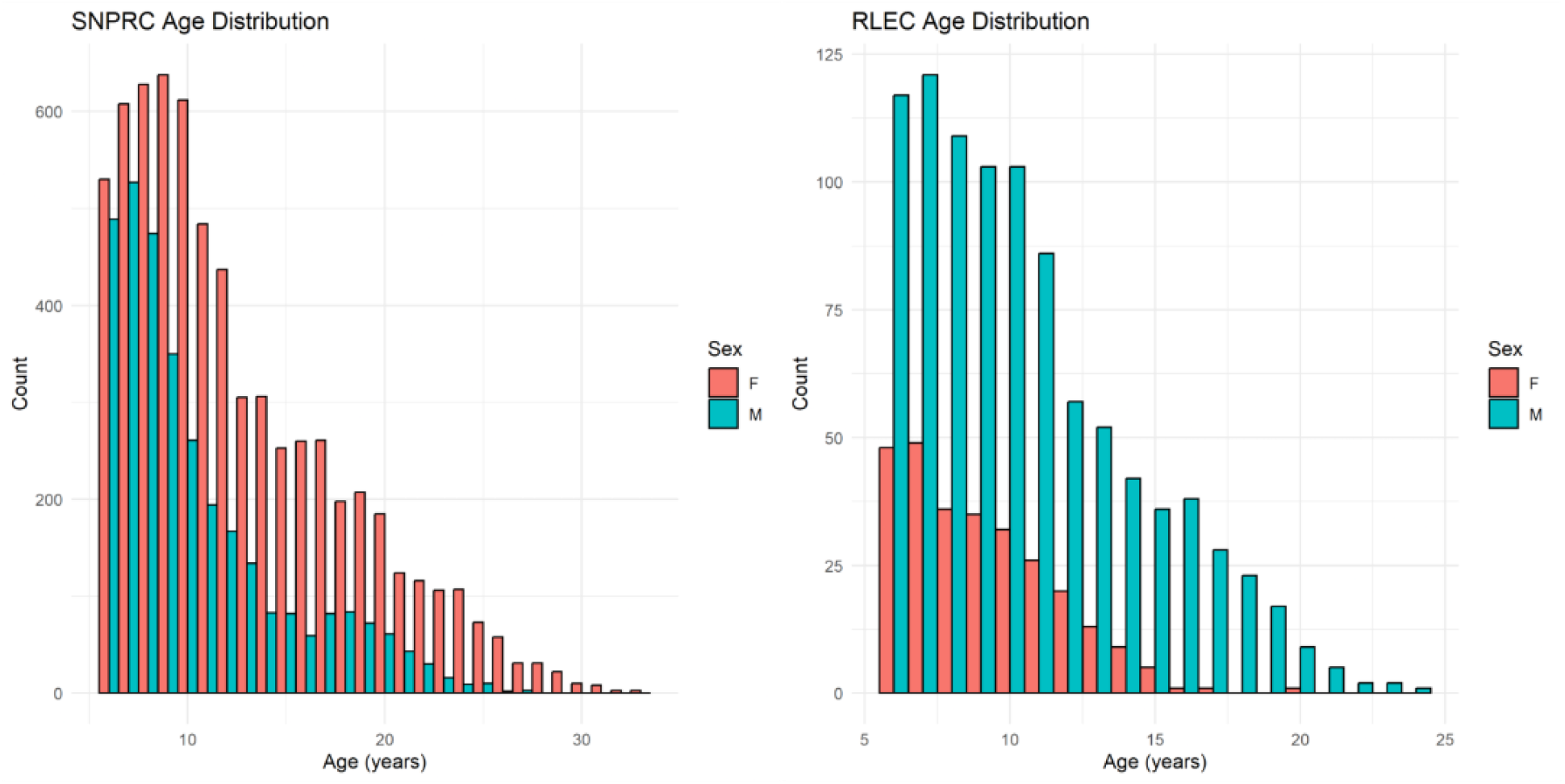
Sex specific distributions of age at observation in the SNPRC and RLEC data sets.

**Figure 2.**
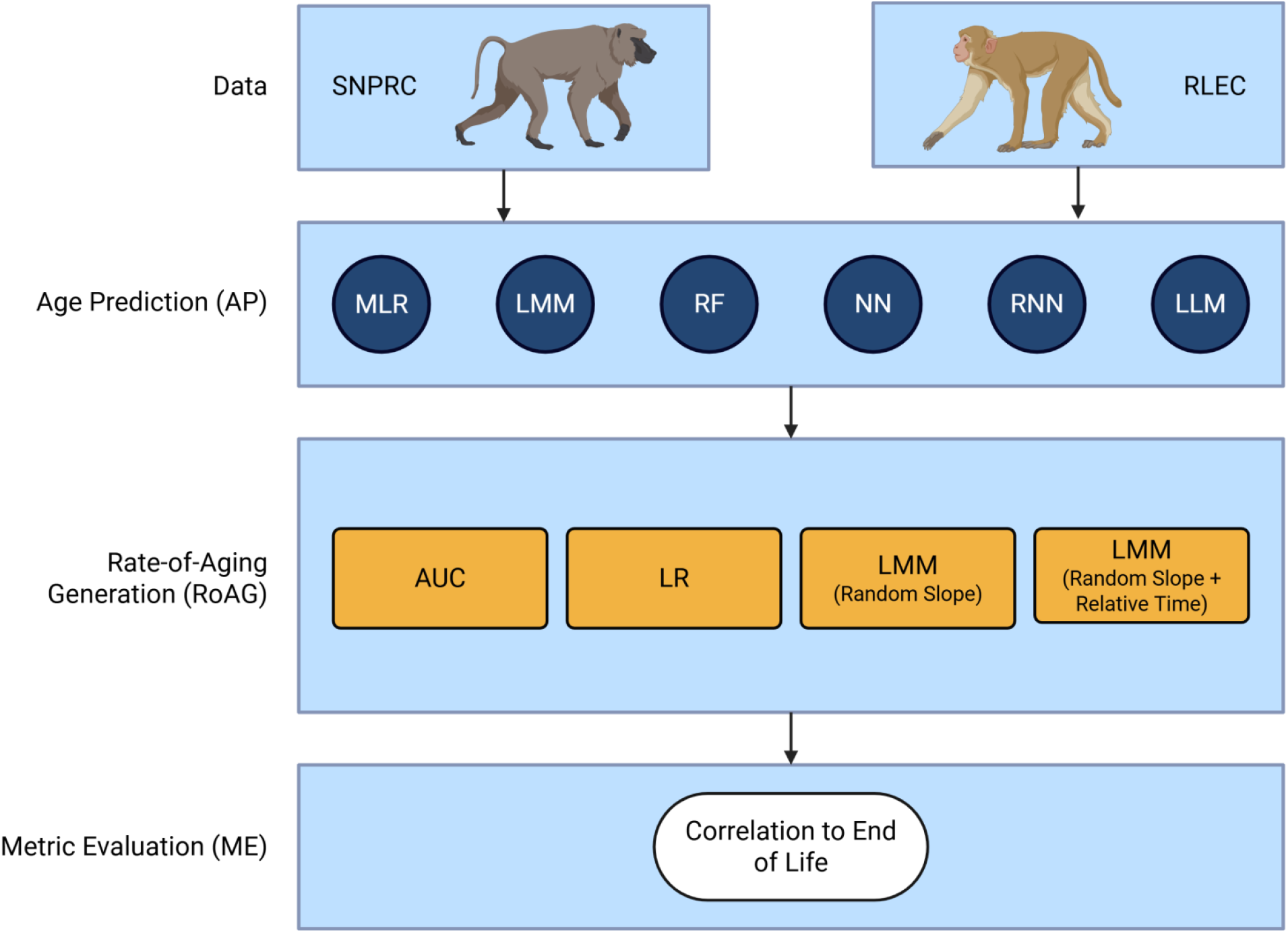
Overview of the pipeline used in this paper, broken into 3 general categories. The acronyms are as follows: MLR – Multiple Linear Regression, LMM – Linear Mixed-effect Model, RF – Random Forest, NN – Neural Network, RNN – Recurrent Neural Network, LLM – Large Language Model, AUC – Area Under the Curve, and LR – Linear Regression.

**Figure 3.**
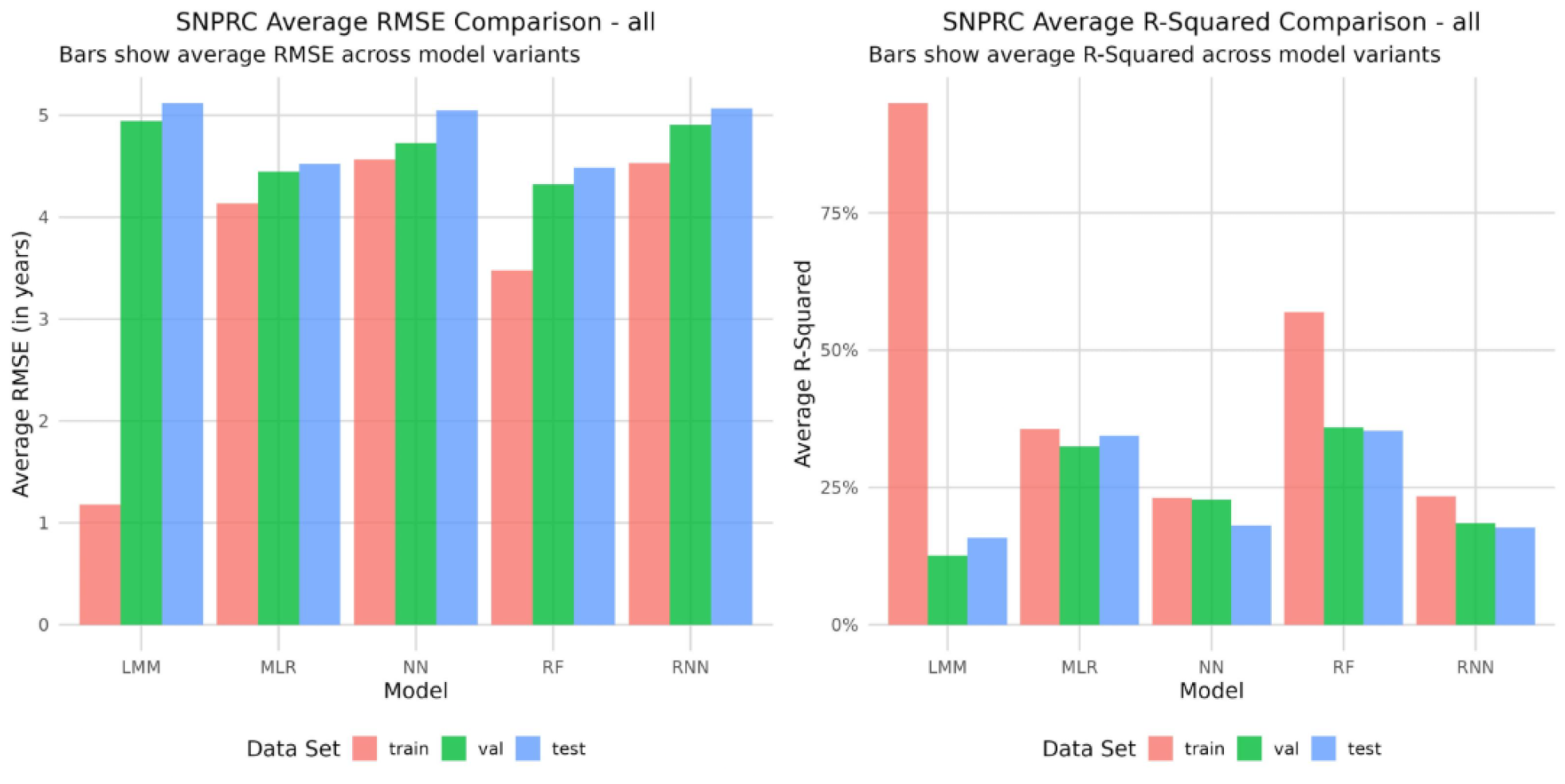
The left side shows the distribution of RMSE and right shows the distributions of R-squared values, across models, and between Train and Test sets, in the SNPRC data.

**Figure 4.**
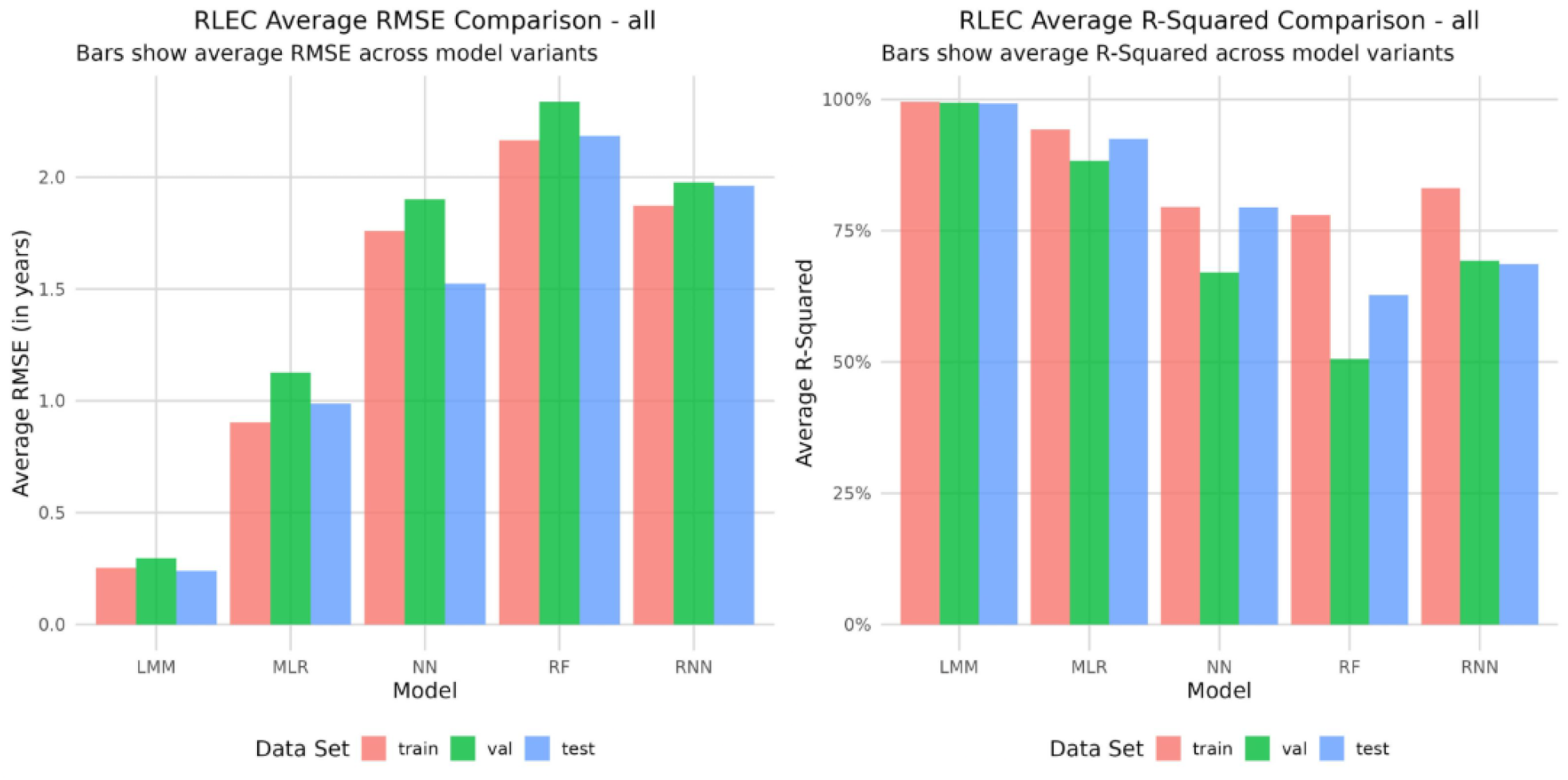
The left side shows the distribution of RMSE and right shows the distributions of R-squared values, across models, and between Train and Test sets, in the RLEC data.

Model training and internal validation were standardized using the rsample (v1.3.0)^13^ and yardstick (v1.3.2)^14^ packages for data partitioning and performance metric calculation. Specific model architectures were implemented using established libraries: Penalized Multiple Linear Regression (MLR) utilized glmnet (v4.1-8)^15^, and Linear Mixed-Effects Models (LMM) were fitted using lme4 (v1.1-36)^16(p4)^ with model parameters extracted via the parameters package (v0.24.2)^17^. The Random Forest (RF) models were constructed using the fast C++ implementation in ranger (v0.17.0)^18^. The deep learning models (NN and RNN) were built and trained using the keras (v2.13.0)^19^ and tensorflow (v2.16.0)^20^ interfaces, with feature importance for the neural networks calculated using the iml package (v1.5.0)^21^. Additionally, an exploratory analysis was conducted using a Large Language Model (LLM), Claude 3 Opus (Anthropic), to assess its regression capabilities.

All code for these analyses can be found at https://github.com/Quillen-Lab/NHPAgingResilience.

### AI-Assisted Technology Disclosure

During the preparation of this work, the authors used Gemini (Google LLC) to assist with code optimization and drafting the text; the authors reviewed and edited the content as needed and take full responsibility for the content of the publication.

### Data Cleaning and Imputation

We did not implement feature selection or reduction based on association with chronological age for this study because the primary focus of our work was to develop a prediction model for biological age, whereas interpretation of associations between predictors and biologic age were of secondary interest. Due to this, we implemented all available features from both datasets that passed filtering and imputation, simplifying the pipeline and avoiding the mistaken removal of informative features. Aside from demographic identifiers (ID, Age, Year, and Sex), the only overlapping features between the datasets were body weight and three specific hematological markers: Hematocrit, Hemoglobin, and White Blood Cell count. Full feature lists are available in the Supplementary Material.

The data processing workflows were implemented in R, using the tidyverse for manipulation and MICE (Multivariate Imputation by Chained Equations)^22^ for imputation. Raw data from both cohorts were aggregated by animal ID, year, age, and sex. Observations within the same year were collapsed by calculating the mean value for each measurement, reducing the data to a single observation per animal per year. Due to this, we did not explicitly account for kinship in the population, since a kinship matrix was not a feasible addition to the data for most models.

Missing values were addressed through a multi-step filtering and imputation process. First, features (columns) and observations (rows) with 75% or more missing data were removed from the dataset because including variables with high missing rates may introduce bias into the analysis. Additionally, for the SNPRC dataset, columns with zero variance or high collinearity (r > 0.98) were identified and removed prior to imputation to prevent numerical instability. The remaining missing values were imputed using the mice package. We employed predictive mean matching for all biological variables, generating 5 imputed datasets with a maximum of 20–50 iterations to ensure convergence.

Post-imputation, specific features in the RLEC dataset still contained a high proportion of missing data or were redundant and were removed. Finally, all continuous biological features were normalized to a mean of 0 and a standard deviation of 1 to standardize the scales for machine learning algorithms, while preserving the original scale of the age variable.

### Data Preprocessing and Partitioning

For both datasets, subjects were filtered to include only animals of known sex and at least 2 observations at ages 6+. To enable the Recurrent Neural Network (RNN) to model temporal dependencies, two temporal features were engineered: Sequence Index representing the ordinal rank of an observation for a subject (e.g., 1, 2, 3), and Sequence Relative Time representing the time elapsed since the subject’s first observation (e.g., 0, 4, 10 years). These features were implemented across all models to maintain consistency.

The data were partitioned into three distinct sets stratified by sex and grouped by individual ID: a training set (70%), a validation set (20%), and a held-out testing set (10%). This ensured that all observations from a single animal remained within the same partition to prevent data leakage. For each dataset, the entire analysis was performed independently on three subsets: all subjects, males only, and females only.

### Model Training and Hyperparameter Tuning

A systematic hyperparameter tuning process was conducted for each model using predefined grids unique to the model’s architecture. Model selection was performed using a multi-criteria Borda ranking method^23^ applied to the training and validation performance metrics. Models were first filtered to exclude those with signs of overfitting (rejecting the top 80% of models with the largest Train-Validation RMSE gaps) and poor accuracy (rejecting the bottom 50% based on Validation RMSE). The remaining models were ranked based on a combination of Training RMSE, Validation RMSE, Training R^2^, Validation R^2^, and the stability of the RMSE gap. The single best-performing model configuration was then evaluated on the final 10% testing set to generate the reported performance metrics.

### Model-Specific Implementation

#### Multiple Linear Regression (MLR)

Penalized regression was implemented using the glmnet package. The hyperparameter grid included alpha values ranging from 0 to 1 (step size 0.5) to test Ridge, Elastic Net, and LASSO penalties. The optimal lambda was determined via an internal 10-fold cross-validation on the training set.

#### Random Forest (RF)

The ranger package was used. Hyperparameters tuned included mtry (5, 6, 7), min.node.size (20, 30, 40), sample.fraction (0.4, 0.5, 0.6), and num.trees (100, 200, 300).

#### Linear Mixed-Effects Model (LMM)

LMMs were fitted using the lme4 package. Two specific model structures were evaluated to handle repeated measures: (1) Random Intercept: Modeled with (1 | ID), including Sequence_idx (visit number) as a fixed covariate; (2) Random Slope for Time: Modeled with (1 + Sequence_relTime | ID), where Sequence_relTime allows for individual aging trajectories but is excluded from the fixed effects to prevent data leakage. Models were fitted using bobyqa and Nelder_Mead optimizers with extended function evaluations (maxfun = 2e5) to ensure convergence.

#### Feed-Forward Neural Network (NN)

NNs were built with keras and tensorflow. The Sex variable was removed during preprocessing to prevent zero-variance issues in sex-stratified subsets. The architecture consisted of two hidden dense layers and a single-unit output layer. An extensive hyperparameter search included the number of units (16–64), dropout rate (0.4–0.6), learning rate (10^-2^–10^-4^, L2 regularization (0.001–0.01), and activation functions (relu, elu).

#### Recurrent Neural Network (RNN)

RNNs were implemented to explicitly model sequential data. Data was reshaped into 3D arrays (Samples x TimeSteps x Features) and padded to the maximum sequence length; a masking layer was utilized to ignore padded values. The architecture consisted of two stacked recurrent layers (GRU or LSTM) followed by TimeDistributed dense output layers, allowing the model to predict age at every time step. Sample weights were applied to mask the loss function for padded time-points. Hyperparameters matched the NN grid, with the addition of cell type (gru, lstm).

#### Large Language Model (LLM)

An exploratory analysis was also conducted using a large language model (Claude 3 Opus) based on a previous publication demonstrating the feasibility of LLMs for regression tasks ^24^. The model was prompted to perform the same age prediction task after a fine-tuning step on the training data. However, this approach yielded extremely poor predictive performance on the test set. Furthermore, the fine-tuning and testing processes incurred significant financial costs related to token usage across all the timepoints. Due to these substantial performance and cost-related hurdles, the LLM was deemed unsuitable for this analytical task and was excluded from the final comparative results.

### Cross-Model Internal Validation

A standardized evaluation pipeline was executed to systematically compare the five modeling approaches. We aggregated the performance metrics from the single best variant of each model type (MLR, LMM, RF, NN, RNN) to assess relative performance. Comparative visualizations included bar plots for Root Mean Squared Error (RMSE) and R^2^ across Training, Validation, and Testing sets, as well as violin plots of prediction residuals to inspect the distribution of errors across the age span.

To determine the most biologically relevant clinical features across the different modeling approaches, we synthesized the feature importance rankings. Since we used internal ranking metrics where possible and different models use different metrics for importance (e.g., coefficients for MLR vs. SHAP values for NN), we converted raw importance scores into ordinal ranks (1–10). We then calculated the Average Rank and Standard Deviation of Rank for each feature across all five models. Features were visualized in a heatmap sorted by Average Rank, highlighting those that were consistently identified as top predictors of biological age regardless of the computational architecture.

### Aging Resilience (AR) Analysis

We derived a set of Aging Resilience (AR) metrics to quantify individual differences in the aging process. These metrics were calculated using prediction residuals (Predicted Age - Chronological Age) generated by the best-performing models. A positive residual suggests accelerated biological aging (lower resilience), while a negative residual suggests decelerated aging (higher resilience). To capture distinct aspects of this trajectory, we calculated four specific metrics classified into two categories: (1) Normalized Cumulative Aging (NCA): This metric quantifies the average burden of “aging error” an individual carries over time. It is calculated as the Area Under the Curve (AUC) of the residuals, normalized by the duration of the observation period (AUC_Norm). This approach treats aging deviation as a cumulative load rather than a momentary rate: (2) Rate of Aging (RoA): These metrics estimate the velocity or acceleration of the aging phenotype. We implemented three variations: (1) RoA-Linear: The slope of a simple linear regression of residuals over chronological age (LM_Slope); (2) RoA-LMM: The individual random slope extracted from a Linear Mixed-Effects Model fitted to the residuals (LMM_Slope); (3) RoA-LMM-Year: A variant of the LMM slope that explicitly controls for the calendar year to account for environmental or colony-management differences over time (LMM_Slope_Year).

### Validation and Robustness Testing

To validate these metrics, we correlated them with the known lifespan of subjects. This analysis was restricted to a “Natural Death” cohort, defined by specific cause-of-death codes indicating euthanasia due to severe age-related decline or serious disease, or due to natural causes. We calculated the Pearson correlation between each AR metric and the animal’s final Age at Death.

Finally, to assess the predictive utility of these metrics earlier in life, we performed a Temporal Cutoff Analysis. The AR metrics were recalculated using only data available up to specific age thresholds (starting from ∼30% of the maximum lifespan and increasing in 3-year increments). This allowed us to determine the earliest point in the lifespan at which the AR metrics became significantly predictive of future mortality.

## Results

### Model Performance Evaluation

We evaluated five approaches (MLR, LMM, RF, NN, RNN) for their ability to predict chronological age. Performance metrics varied significantly between the two cohorts, reflecting differences in feature types and dataset structures.

In the SNPRC cohort, overall predictive power was moderate. The Random Forest (RF) and Multiple Linear Regression (MLR) models achieved the highest accuracy, with Test R^2^ values of 0.353 and 0.344, respectively. The Linear Mixed-Effects Model (LMM), Neural Network (NN), and Recurrent Neural Network (RNN) generally yielded lower predictive accuracy for chronological age (R^2^ < 0.20), highlighting the difficulty of modeling age solely from standard CBC and blood chemistry data in this population.

The RLEC cohort analysis yielded high predictive accuracy. The LMM and MLR models emerged as the top performers for age prediction. The LMM achieved a near-perfect Test R^2^ of 0.992, while the MLR achieved 0.925. In comparison, the non-linear models (NN and RNN) showed lower performance (R^2^ of 0.794 and 0.686, respectively).

### Biological Feature Importance

To identify the biological drivers of the predicted aging trajectory, we synthesized feature importance rankings across all five models (Figure 5). In SNPRC baboons, the most consistent biological predictors of age were related to hematological and immunological function in line with features that predominate in the input. Beyond generalized changes in body weight, the models prioritized Red Blood Cells and Mean Platelet Volume. This suggests that in the baboon cohort, the measurable aging process is phenotypically dominated by declining circulatory efficiency and “inflammaging”—a state of chronic, low-grade systemic inflammation^25^. However, it is important to note that blood-based biomarkers dominated the input features for the SNPRC baboons. For the RLEC macaques, where a more diverse range of features was available, the aging signature in this cohort was cardiometabolic. The consensus top features included Whole Body Mass, Alkaline Phosphatase (Alp_U.L), and cardiac structural metrics such as Left Ventricular Internal Diameter (LVID). This indicates that for the rhesus macaques in this specific cohort, structural changes in the heart and shifts in metabolic homeostasis are the primary signals of biological aging captured by the models. It is important to note that these datasets have drastically different feature sets, which significantly contributed to the differences in feature importance.

**Figure 5.**
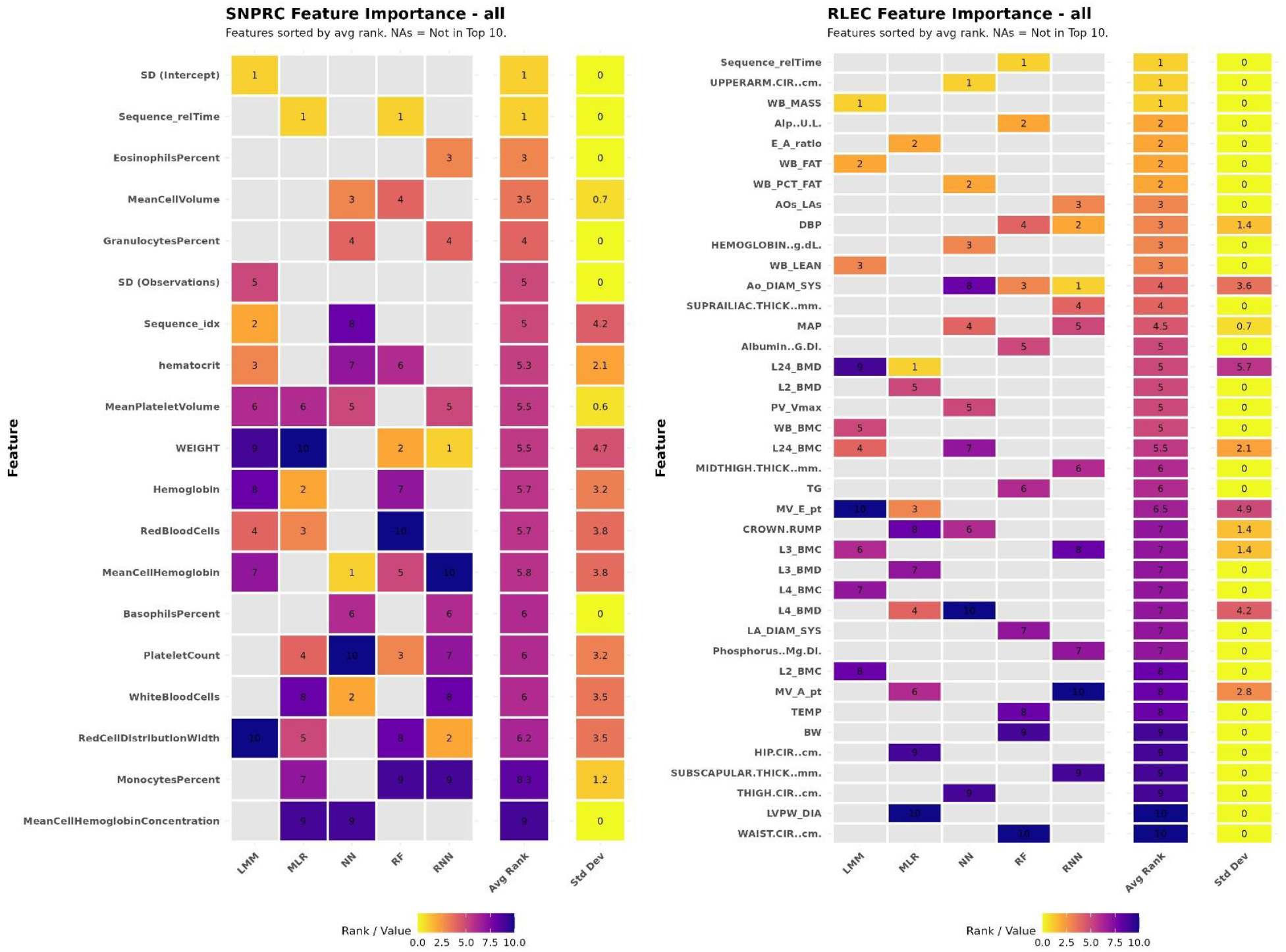
This figure shows the top 10 features across all the models and datasets. The rows are ranked by Avg Rank, which is the sum of the rank number for each feature, divided by its number of occurrences.

### Sex-Specific Aging Dynamics

Sex-stratified analyses revealed notable differences in model performance. In the SNPRC cohort, models trained on male subjects generally achieved higher accuracy than those trained on females. For example, the RF model achieved a Test R^2^ of 0.512 in males compared to 0.328 in females. Feature importance analysis suggested distinct aging drivers: Hemoglobin levels were the strongest predictor for males, emphasizing oxygen-carrying capacity, whereas inflammatory markers were more predictive for females. In the RLEC cohort, performance remained high across sexes, though males (LMM R^2^ = 0.996) were predicted with slightly higher precision than females (LMM R^2^ = 0.978). Alkaline Phosphatase emerged as a consistent, sex-independent marker of aging in the macaques, ranking highly for both males and females.

### Validation of Aging Resilience (AR) Metrics

We derived Aging Resilience (AR) metrics—specifically Normalized Cumulative Aging (NCA) and Rate of Aging (RoA)—from the model residuals and validated them against the known lifespan of animals in the “Natural Death” cohort.

Correlation analysis (Figure 6) demonstrated that while linear models were superior at predicting chronological age, the AR metrics derived from neural network architectures (RNN and NN) were the strongest predictors of mortality. In the RLEC cohort, despite the LMM’s near-perfect age prediction, the NCA derived from the RF model yielded the highest correlation with Age at Death (r > 0.8). In the SNPRC cohort, the RNN-derived NCA showed a strong correlation with lifespan (r > 0.8). This confirms that while physical characteristics (captured by linear models) track time well, the non-linear dynamics captured by the RNN and RF better reflect the biological deterioration that limits lifespan. The RoA (velocity-based) metrics underperformed, losing to the NCA (cumulative load) metrics in predicting lifespan, suggesting that the rate of physiological decline is a less acute (or more difficult to capture) indicator of mortality risk than the total accumulated deficit burden in these models.

**Figure 6.**
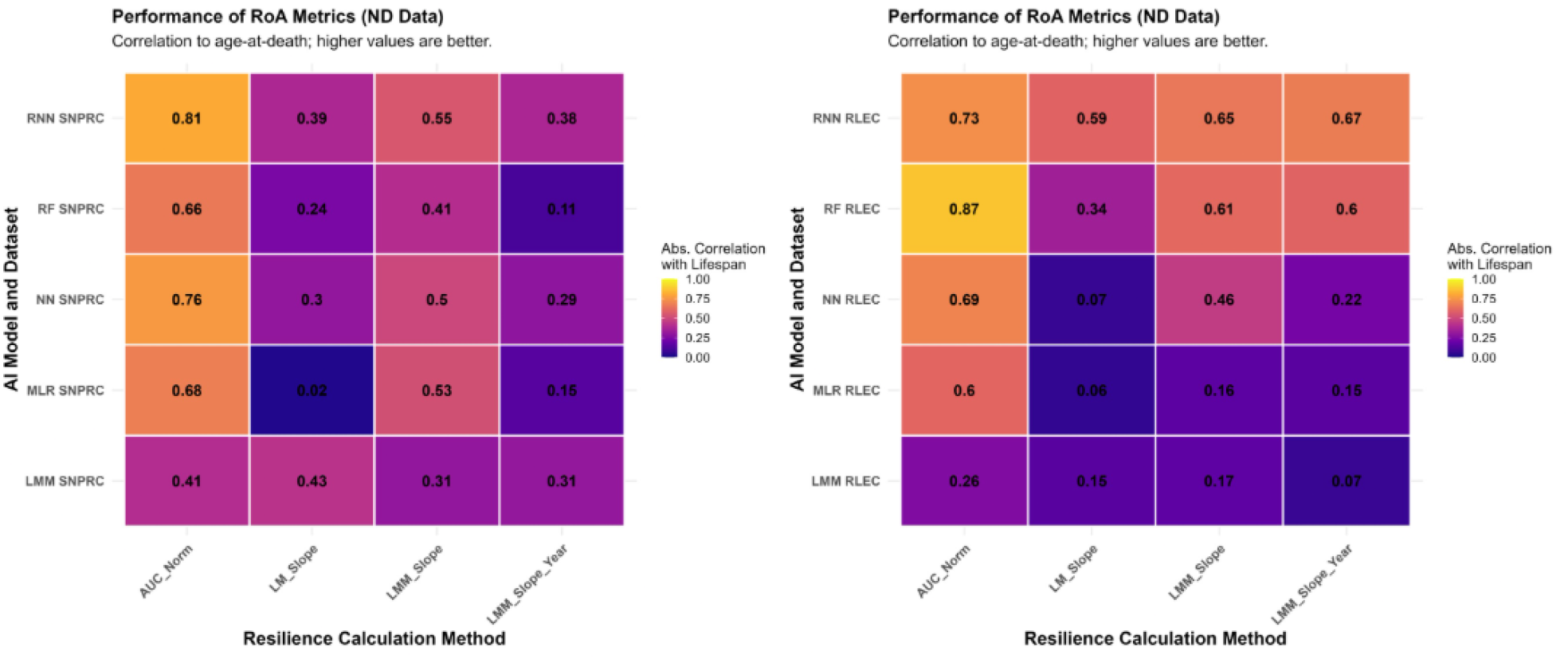
Left is SNPRC and right is RLEC; This figure shows the distribution of correlations between model combinations for each AR metric.

### Temporal Robustness

To assess the clinical utility of these metrics for early prediction, we performed a temporal cutoff analysis (Figure 7). We recalculated the AR metrics using data truncated at 3-year incremental age thresholds (e.g., using only data up to 9 years, 12 years, etc.). Unfortunately, the AR metrics could not capture meaningful correlations to Age at Death, until data began approaching late-life metrics (as seen above in Figure 7, depicted cutoff analyses at ages 9 and 21). Even then, the most correlated AR metrics were derived from the model that massively overfit the data during the Age Prediction step and only performed well with data leading later into life. While there were some standout correlations at each stage, no model/metric combination consistently performed well across cutoffs. The RLEC data also failed in multiple steps, due to lack of data necessary to create the metrics and calculate their correlations, which is why it is not present at the “21_years” plot above.

**Figure 7.**
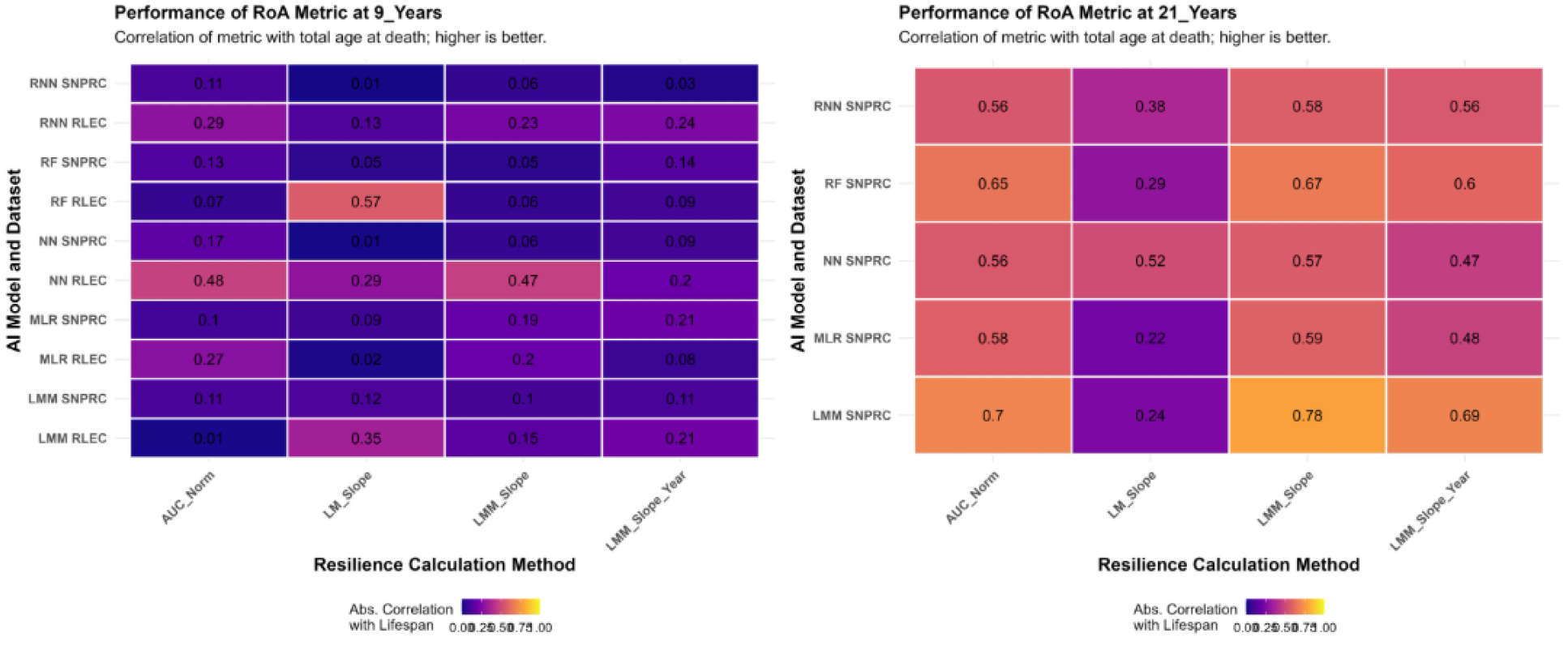
This figure shows the distribution of correlations between model combinations for each AR metric, using the lowest and highest cutoff metrics for each data set.

## Discussion

This study’s most significant contribution is the validation of an Aging Resilience (AR) metric capable of strongly predicting lifespan from routine clinical data. While predicting chronological age is a useful intermediate step, the ability to forecast mortality risk in living animals provides a far more powerful and translationally relevant tool for geroscience. Our findings that the RNN and RF derived ARs (particularly, the NCA) correlated strongly with lifespan in their respective cohorts (Pearson’s r > 0.8) provides compelling evidence that the dynamics of physiological aging, rather than a single static age prediction, are a key indicator of healthspan. This is particularly valuable for NHP research, where animals are often humanely euthanized at the first sign of serious illness, making the AR metric a more useful one than traditional frailty indices that are primarily or exclusively focused on identifying older adults with low resilience (i.e. frail vs. non-frail), rather capturing variability across the lifespan.

The NCA metric quantifies an individual’s average burden of aging, where a higher value represents a greater burden relative to chronological age. Biologically, the models capture this deterioration through shifts in key clinical markers. For example, a high AR in the SNPRC cohort may reflect worsening hematological profiles and chronic inflammation, while in the RLEC cohort, it could signify a rapid decline in cardiopulmonary function, given our feature importance results. Therefore, AR metrics serves as an integrated measure of this multi-system decline. This is crucial given that clinical data sets are heterogeneous. A higher AR value indicates that an animal is more-readily accumulating this age-related damage, regardless of the specific system involved, which is consistent with our finding that this metric strongly predicts an earlier end of life.

Our systematic comparison of five distinct modeling approaches revealed that non-linear models (specifically RNN and RF) consistently provided the best balance between predictive accuracy, generalization to unseen data, and ability to capture biological relevance of aging (as seen in AR results). While linear models (MLR and LMM) achieved exceptionally high accuracy in predicting chronological age in the RLEC cohort, this was likely driven by the inclusion of physical characteristic features, such as DEXA scan values (e.g., bone mineral content, whole body soft tissue mass), which were not available in the SNPRC dataset. These features often exhibit strong, linear declines across the lifespan that track closely with chronological time, allowing linear models to map these physical traits to age with high precision. Furthermore, RLEC animals are highly heterogeneous in their exposure to physiological stressors with most having undergone radiation exposure and all being maintained on a Typical American Diet. The SNPRC cohort similarly contains animals having undergone a variety of short-term diet challenges, but no pervasive challenges to early-life resilience. However, our validation analysis revealed a critical paradox: while these models were nearly perfect at predicting *how old* an animal was, they were poor at predicting *when they would die* (low correlation with lifespan). This discrepancy highlights a fundamental distinction between “chronological aging”—the linear passage of time marked by gross physical changes—and “biological resilience,” which is dynamic and non-linear. The non-linear models, though less accurate at pinpointing exact chronological age, were significantly better at predicting mortality, suggesting they captured the complex, non-linear perturbations in homeostatic maintenance (e.g., inflammation, metabolic shifts) that precipitate death.

We note the stark difference in chronological age prediction performance between the two cohorts. The substantially higher variance captured, and accuracy achieved in the RLEC dataset (Test R^2^ of ∼0.3-0.9 and Test RMSE of ∼0.2-2.1) compared to the SNPRC dataset (Test R^2^ ∼0.1-0.5 and Test RMSE of ∼4.5-5.1) is likely driven by the difference in feature depth (80 in RLEC vs. 19 in SNPRC) and type. The RLEC data included specialized cardiac and metabolic data which track development and aging closely. However, despite these differences in initial age prediction accuracy, the AR metrics derived from both cohorts showed strong predictive validity for mortality. This discrepancy emphasizes that a “good” aging clock is not necessarily one that predicts chronological age perfectly, but one that captures the biologically relevant variance associated with whole-system health outcomes.

The sex-stratified analyses also revealed important differences in aging dynamics. In the SNPRC cohort, where sample sizes were sufficient for a meaningful comparison, models were more accurate for males. Meanwhile, in the RLEC, the inverse is true, outside of the RNN (likely due to large sample size differences between sexes). These differences were expected, given the sexual dimorphism of these species, and were included to verify those assumptions.

These findings position the AR metrics as a significant advancement over existing NHP frailty assessments. While previous work pioneered establishing the “frail phenotype” in rhesus macaques^8^, it relied on a deficit accumulation model that requires overt clinical decline, which is not always applicable for NHP cohorts. In contrast, our AI-derived metrics function analogously to the “aging clocks” recently developed for mice^9^, but with the critical advantage of being implemented in a translational primate model. By shifting from a classification of frailty to a continuous measure of resilience, our approach allows for the detection of subtle biological deviations in the “healthy” phase of lifespan, potentially filling the translational gap between murine molecular clocks and human clinical frailty indices. Critically, because this approach utilizes standard clinical values found in human Electronic Health Records (EHR), this methodology could be directly adapted to monitor patient resilience and validate interventions in human healthcare settings without the need for novel biomarkers.

A primary strength of this study lies in the use of two independent, highly distinct cohorts to validate the methodology. By testing our models on both the large, sparse SNPRC dataset (baboons) and the smaller, feature-rich RLEC dataset (macaques), we demonstrated that the “Aging Resilience” concept is robust to differences in species, sample size, and data dimensionality. Furthermore, unlike many aging clock studies that rely solely on chronological age as a ground truth, we validated our metrics against a hard biological endpoint: natural death. This validation against the “Natural Death” portion of each cohort ensures that our metrics are not merely tracking the passage of time, but are capturing the actual biological attrition that limits lifespan.

This study has several limitations. First, while we can identify which features are most important in the NN and RNN models, their “black box” nature complicates direct biological interpretation. For example, we cannot easily determine if a higher or lower red blood cell count is driving the prediction of accelerated aging without further targeted analysis, due to SHAP not providing direction of effect for its importance ranking. Second, while non-human primates are excellent models for human aging, the direct translation of these specific aging clocks to human populations requires further investigation and external validation. Finally, our AR calculations, while effective, represent only a few potential methods to quantify aging dynamics; future work could explore more robust calculations that may be more biologically informative.

In conclusion, this work establishes a robust methodology for developing and validating aging clocks from routinely collected repeated measures clinical data. While the specific clocks generated here are cohort-specific, the methodology is translatable. We recommend that future studies seeking to develop similar clocks prioritize collecting a wide array of deeply phenotyped, longitudinal data. Most importantly, we show that the resulting AR metric is a strong predictor of healthspan, providing a valuable tool for monitoring the health of living animals and for quantitatively assessing the impact of potential anti-aging interventions in preclinical NHP trials.

## Acknowledgements

This work was supported by the National Institute on Aging at the National Institutes of Health (grant number R01 AG087957 to E.E.Q.); the National Institute of Allergy and Infectious Diseases at the National Institutes of Health (grant number U01 AI150578 to J.M.C.); and the Office of Research Infrastructure Programs at the National Institutes of Health (grant number P51 OD011133 to SNPRC). We thank Deborah Newman and Terry Hawkins for their tremendous effort in compiling and providing the SNPRC data set as well as the many veterinarians, technicians, and others who generated this data over many decades.

