## Supplementary Material for "Developing And Internally Validating AI-Based Aging Resilience Biomarkers in Non-Human Primates"

### **Supplementary Materials**

SNPRC Features – ID, Year, Age, Sex, WEIGHT, BasophilsPercent, EosinophilsPercent, GranulocytesPercent, hematocrit, Hemoglobin, LymphocytesPercent, MeanCellHemoglobin, MeanCellHemoglobinConcentration, MeanCellVolume, MeanPlateletVolume, MonocytesPercent, PlateletCount, RedBloodCells, RedCellDistributionWidth, WhiteBloodCells

RLEC Features – Age, ID, Sex, Year, BUN(mg/dL), Calcium(mg/dL), Creatine(mg/dL), Glucose(mg/dL), Hematocrit(%), Hemoglobin(g/dL), Phosphorus(mg/dL), TSP(g/dL), WBC(10^3/uL), Albumin(g/dL), ALP(U/L), AST(U/L), Chloride(mEq/L), CholesterolTotal(mg/dL), Potassium(mEq/L), Sodium(mEq/L), AnionGap(mEq/L), TCO2(mEq/L), ARToLA(ratio), AortaDiamSyst(mm), Bodyweight(kg), DiastBloodPressure(mmHg), LVEndDiastVolA4C(mL), LVEjectionFraction(%), MVEndPointSeptalSep(cm), LVEndSystVolA4C(mL), FractionalShortening(%), HDLTotal(mg/dL), HeartRate(bpm), Insulin(mU/L), IVSThickDiast(cm), IVSThickSyst(cm), LADiamSyst(mm), LVInternalDiamDiast(cm), LVInternalDiamSyst(cm), LVPostWallThickDiast(cm), LVPostWallThickSyst(cm), MeanArterialPressure(mmHg), RespirationRate(br/min), SystBloodPressure(mmHg), SpO2(%), Temperature(F), Triglyceride(mg/dL), AbdominalThick(mm), AVPeakVel(m/s), ChestThick(mm), Cortisol(ug/dL), Estradiol(pg/mL), MVEARatio, HipCircum(cm), Lumbar24BMC(gm), Lumbar24BMD(gm/cm^2), Lumbar2BMC(gm), Lumbar2BMD(gm/cm^2), Lumbar3BMC(gm), Lumbar3BMD(gm/cm^2), Lumbar4BMC(gm), Lumbar4BMD(gm/cm^2), MidscapularThick(mm), MidthighThick(mm), MVAWavePeakVel(m/s), MVEWavePeakVel(m/s), PVPeakVel(m/s), SubscapularThick(mm), SuprailiacThick(mm), Testosterone(ng/dL), ThighCircum(cm), TricepThick(mm), TrunkLength(cm), UpperarmCircum(cm), WaistCircum(cm), WholeBodyBMC(gm), WholeBodyBMD(gm/cm^2), WholeBodyFat(gm), WholeBodyLean(gm), WholeBodyMass(gm), WholeBodyFat(%), HemoglobinA1C(%), LDLTotal(mg/dL), CrowntoRumpLength(cm), AVPeakGradient(mmHg), AVMeanGradient(mmHg), AVVelTimeInterval(cm), AVMeanVel(m/s), MVAERatio, MVLatAnnulusPeakVelLateDiast, MVSeptAnnulusPeakVelLateDiast, LVCardiacOutput(1/min), LVCardiacOutputA2C(1/min), LVCardiacOutputA4C(1/min), MVDecelerationSlope(m/s^2), MVDecelerationTime(ms), LVMajorAxisLengthDiffD(%), LVMajorAxisLengthDiffS(%), LVEndDiastVol(mL), LVEndDiastVolA2C(mL), EndDiastVolTeich(ml), LVEjectionFractionA2C(%), LVEjectionFractionA4C(%), EjectionFractionTeich(%), LVEndSystVol(mL), LVEndSystVolA2C(mL), EndSystVolTeich(ml), MVEToMVLatAnnulusEaRatio, MVLatAnnulusEaToAaRatio, MVSeptAnnulusEaToAaRatio, MVLatAnnulusPeakVelEarlyDiast, MVSeptAnnulusPeakVelEarlyDiast, LVGlobalLongitudinalStrain(%), LAAreaSystA2C(cm^2), LAAreaSystA4C(cm^2), LAEndSystVolA2C(ml), LAEndSystVolA4C(ml), LAEndSystVolBP(ml), LAToARRatio, LVOutflowTractDiam(mm), PVPeakGradient(mmHg), LVStrokeVol(ml), LVStrokeVolA2C(ml), LVStrokeVolA4C(ml), StrokeVolTeich(ml)

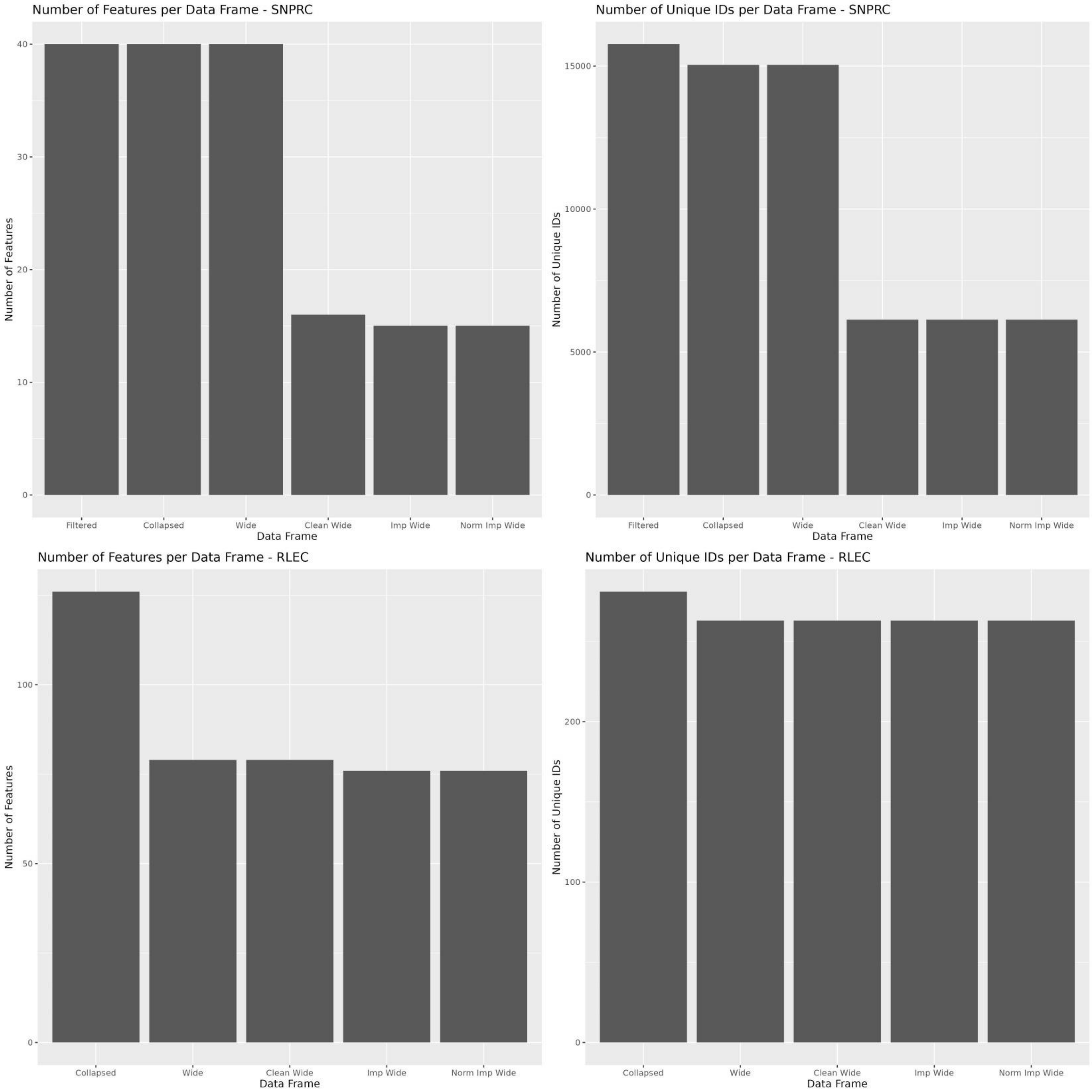


The above plots represent the number of unique IDs and features that were lost as data was cleaned, transformed, and imputed.

1. Filtered (SNPRC only) – Standardized feature naming and averaged ID/Day/Measurement
2. Collapsed – ID/Year/Measurement combinations were averaged, so there was only one ID/Year/Measurement combination per ID.
3. Wide – Converted long format data into wide format, so that all the features represented in Measurement get their own columns.
4. Clean Wide – Removed rows that had over 75% missing data, then removed features missing over 75% of its data, 0/NA variance, and any that are highly colinear.
5. Imp Wide – Imputed missing values and removed any features that were unable to be fully imputed.
6. Norm Imp Wide – Implemented sex specific z-score normalization across all the numeric features, aside from ID, Year, and Age.
